# Microbiome-Dependent Protection Against *Corynebacterium bovis*-Associated Hyperkeratosis in Nude Mice (*Mus musculus*)

**DOI:** 10.64898/2026.04.11.716586

**Authors:** Kate E. Fodor, Amanda C. Ritter, Rebecca A. Schmieley, Ileana C. Miranda, Rodolfo J. Ricart Arbona, Neil S. Lipman

## Abstract

*Corynebacterium bovis*, the cause of *Corynebacterium*-associated hyperkeratosis (CAH), is an important pathogen in immunocompromised mice that is difficult to eliminate and can confound research outcomes. We recently observed that CAH severity varies among outbred athymic nude mouse stocks, but the relative contributions of host genetics and the microbiome remain unclear. We hypothesized that disease course and severity vary based on host genetic stock and/or microbiome composition. Three nude mouse stocks were rederived into the axenic state and either monoinfected with a pathogenic *C. bovis* isolate (10^4^; CFU) or given sterile media (n=6/group). Axenic mice were also reassociated with their source microbiome or microbiomes from three other stocks with known differences in CAH severity, then inoculated with *C. bovis* (n=6) or sterile media (n=2). In a separate experiment, one axenic stock was used to assess the role of *C. amycolatum* via monoinfection, monoinfection followed by *C. bovis* challenge, or addition to a nonprotective microbiome followed by *C. bovis* challenge. Mice were monitored daily for 21 days and scored for skin lesions (0-5). *C. bovis* monoinfected mice developed disease comparable in severity and timing to conventionally raised controls. Notably, reassociation with Vendor A2’s microbiome prevented clinical lesions and reduced histopathologic changes across all stocks. While *C. amycolatum* as a monoinfection did not cause disease nor reduce disease severity following *C. bovis* challenge, it delayed the onset and lowered peak scores when added to a non-protective microbiome. These findings demonstrate that *C. bovis* can cause CAH as a monoinfection, that both host genetics and microbiome composition influence disease progression, and, together with prior work, support its role as the etiologic agent consistent with Koch’s postulates. Identifying protective microbiome constituents may inform strategies to reduce disease burden in susceptible mice.

## Introduction

*Corynebacterium*-associated hyperkeratosis (CAH), or scaly skin disease, is a clinically significant condition of immunocompromised mice caused by the opportunistic pathogen *Corynebacterium bovis*.^1,2^ Affected mice commonly present with hyperkeratotic dermatitis, acanthosis, erythema, alopecia in hirsute strains, weight loss, dehydration, pruritus, and a hunched posture.^1,3-5^ Once established within a colony, *C. bovis* is difficult to eliminate because naïve mice are readily colonized through contaminated fomites or skin effluvium by airborne and fecal-oral routes.^6,7^ Although select antibiotics can reduce clinical signs, none eradicate infection,^8-10^ and treatment can alter the microbiome or induce lethal *Clostridioides difficile* disease in highly immunocompromised strains.^11^ Uncontrolled *C. bovis* infection may also confound research outcomes by increasing morbidity and reduced engraftment of patient-derived xenografts.^12^

Athymic nude mice typically develop CAH 7-10 days after infection, with lesions resolving over a similar period.^1,2,13^ Disease severity is influenced by inoculum dose, strain, route of infection, and host age.^1,4^ Recent work demonstrated that host stock and microbiome composition are additional determinants of CAH severity; outbred athymic nude mice from different vendors show markedly different clinical outcomes despite receiving identical inocula.^2^ These differences likely reflect both genetic variation among stocks and differences in their associated microbiomes.^14-16^

The skin microbiome of immunocompromised mice frequently includes commensal or pathobiont corynebacteria and staphylococci, including *Corynebacterium amycolatum, Corynebacterium mastitidis*, and *Staphylococcus xylosus*.^9,17,18^ Recent findings by Michelson et al.^2,19^ demonstrated that *C. amycolatum* is a prevalent skin commensal in certain athymic nude mouse stocks and may be associated with reduced CAH severity.^2,19^ In these studies, stocks harboring *C. amycolatum* prior to *C. bovis* exposure exhibited attenuated clinical disease compared with stocks lacking this organism, suggesting a potential protective or competitive role within the skin microbiome.^2,19^ Inconsistent CAH development following controlled *C. bovis* exposure suggests that disease expression is multifactorial.^2,4,20^ Koch’s postulates have not yet confirmed *C. bovis* as a sole etiologic agent, and the contribution of host genetics to susceptibility remains poorly defined. Although CAH is most common in immunodeficient strains, cases have also been reported in immunocompetent mice,^21^ and variation in disease severity among athymic nude stocks suggests that additional genetic factors beyond the *Foxn1* mutation modulate susceptibility.^2^

Given the widespread use of immunocompromised mouse models and the prevalence of *C. bovis* across research institutions, improved strategies to mitigate CAH and limit its impact on research outcomes are needed. A clearer understanding of the interactions among host genetics, microbiome composition, and *C. bovis* virulence is essential for developing effective mitigation strategies. Our laboratory has previously characterized the epizootiology and clinical manifestations of *C. bovis* infection.^1-3,6,8^

The current study aimed to determine whether *C. bovis* can cause CAH as a monoinfection in axenic outbred athymic nude mice and evaluate how host stock and microbiome source influence disease severity. Axenic outbred athymic nude mice from multiple vendors were used to test the following hypotheses: (1) monoinfection with pathogenic *C. bovis* will consistently induce CAH, fulfilling Koch’s postulates; (2) disease severity will differ among stocks, indicating genetic variation in susceptibility; (3) reassociation of axenic mice by cohousing with conventionally raised mice with unique microbiota will prevent CAH or reduce its severity, implicating microbiome constituents in disease modulation; and (4) prior colonization of axenic nude mice with *C. amycolatum* will attenuate CAH severity following subsequent *C. bovis* infection, suggesting a disease-modulating role.

## Materials and Methods

### Experimental Design

Three distinct genetic stocks of athymic nude mice obtained from commercial vendors (Stock A [Hsd:Athymic Nude-*Foxn1nu*], Stock B [J:NU], and Stock C [Crl:NU(NCr)-*Foxn1nu*]) were rederived to the axenic (germfree) state (F0 generation). Equal numbers of male and female homozygous nude mice from subsequent litters (F1 generation) served as experimental animals (Figure 1).

**Figure 1.**
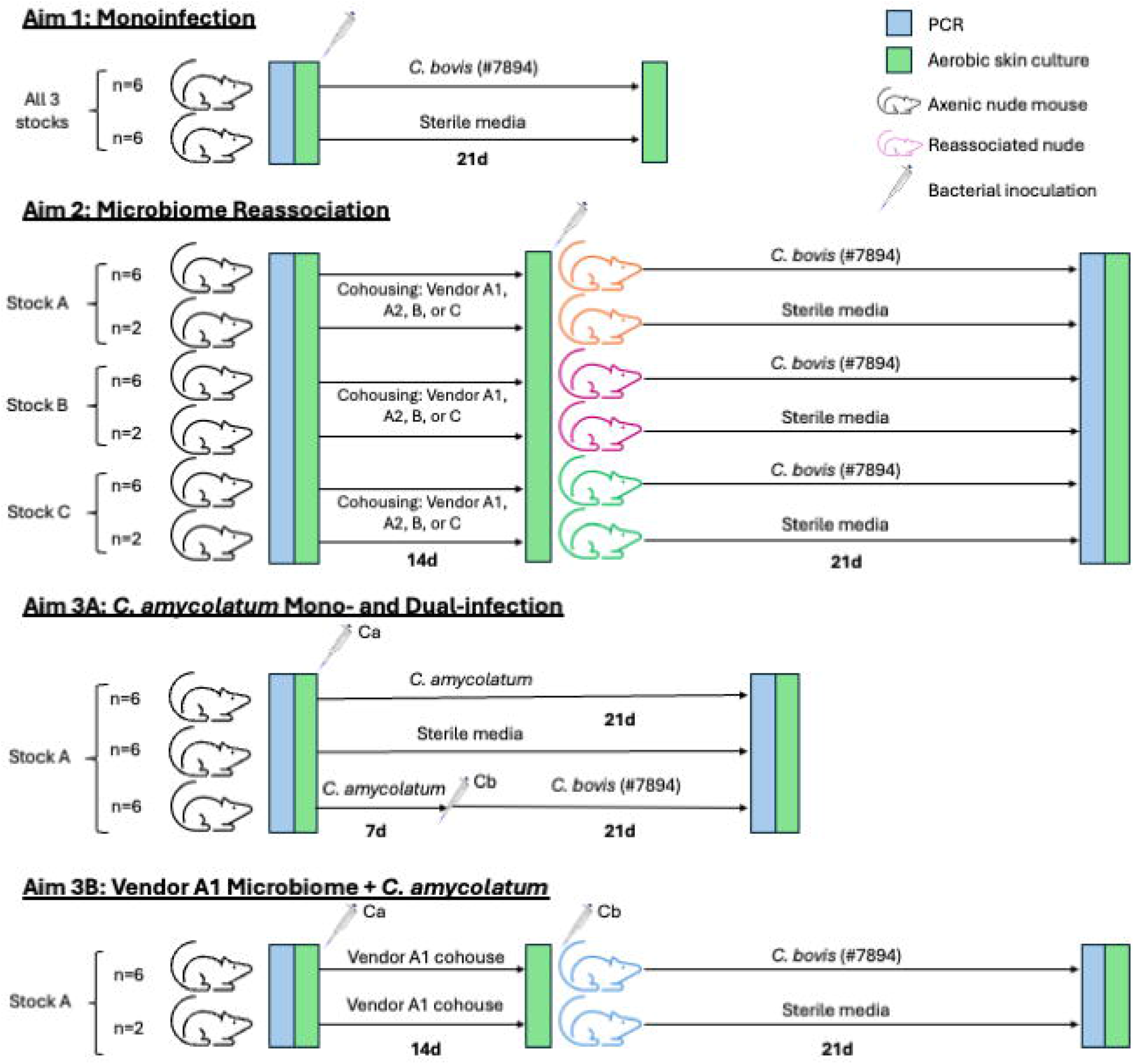
Schematic detailing temporal events and samples collected/assays performed for experiments conducted for each study aim. Mice inoculated with *C. bovis* isolate 7894 received 10^4^ CFU/mouse; mice inoculated with *C. amycolatum* received 10^8^ CFU/mouse; and negative controls received 50 μL of sterile media topically. Aerobic culture samples were collected at multiple timepoints.

#### Aim 1: C. bovis monoinfection

Axenic mice from each stock (n=6) were topically inoculated with 10□ CFU of pathogenic *C. bovis* field isolate 7894, or sterile bacteria-free media (negative control) (Figure 1).^1,2^ Clinical signs were assessed daily and scored 0 to 5 based on lesion severity (Table 1). Mice were euthanized by CO□ overdose at 21 days post-inoculation (dpi) or if clinical signs reached grade 5 (humane endpoint). *C. bovis* colonization was confirmed by aerobic culture before inoculation and at the experimental endpoint. A complete gross necropsy was performed, macroscopic changes documented, and 6 full-thickness skin sections per mouse collected and scored histologically.

**Table 1.** *C. bovis* clinical scoring system for nude mice.

| Grade | Clinical signs |
| --- | --- |
| 0 | No clinical signs observed |
| 1 | Few flakes or scales adhered to the skin or small dermal papules without flaking. |
| 2 | Few flakes or scales adhered to the skin or mildly hunched posture and mild erythema associated with flaking or scaly lesions. |
| 3 | Moderate erythema or moderate number of white flakes adherent to the skin or few white flakes at 3 or more additional locations affected, including the ventrum, muzzle, cheeks, crown, and limbs. |
| 4 | Moderate erythema or moderate number of white flakes adherent to the skin and moderately hunched posture. |
| 5 | Numerous flakes or scales adhered to skin (thick layer) or severe erythema or severely hunched posture. |

Reprinted in part with permission from Mendoza et al.<sup>1</sup>

#### Aim 2: Microbiome reassociation and C. bovis challenge

Axenic mice from each stock were cohoused for 2 weeks with conventionally raised athymic nude mice from one of 4 sites (one from each stock’s originating colony, plus an additional colony from Vendor A where Stock A’s microbiome includes *C. amycolatum*, a species not detected at the vendor’s other location) to enable microbiome reassociation with vendor-specific microbiomes (Figure 1). This process generated 24 unique combinations of stock, microbiome, and sex. After cohousing, the donor mouse was removed, and the experimental mice were topically inoculated with 10□ CFU of the pathogenic *C. bovis* isolate 7894 (n=6) or sterile bacteria-free media (negative control, n=2). Mice were scored daily for cutaneous acanthosis and hyperkeratosis (CAH) lesions until 21 dpi as described for Aim 1. *C. bovis* colonization was assessed prior to inoculation and at experimental endpoint via aerobic culture.

#### Aim 3: Assessment of C. amycolatum’s pathogenicity and protection

Axenic Stock A mice (n=6) were monoinoculated with 10□ CFU of *C. amycolatum* isolate 25-1107-1 and monitored daily for skin lesions for 21 dpi (Figure 1). Additional axenic Stock A mice (n=6) were topically inoculated with *C. amycolatum*, confirmed by aerobic culture to be colonized on 7 dpi, and subsequently inoculated with *C. bovis* isolate 7894 (10^4^) with skin lesions assessed and scored for 21 dpi following *C. bovis* inoculation. Samples were collected at the study endpoint as described for Aim 1. Additionally, axenic Stock A mice were reassociated with Stock A’s microbiome from Vendor A’s *C. amycolatum*-free site and were concurrently inoculated with 10□ CFU *C. amycolatum* when initiating cohousing. The mice were subsequently inoculated with *C. bovis* isolate 7894 two weeks after initiating cohousing, following removal of the microbiome donor, and were assessed daily for 21 dpi post *C. bovis* infection to determine whether adding *C. amycolatum* to this microbiome altered the disease course. Samples from this cohort were collected and assessed as described for Aim 2.

### Animals

Heterozygous female (*Foxn1nu/+*) and homozygous male (*Foxn1nu/nu*) outbred athymic nude mice, 5 to 8 weeks of age, were purchased from 3 commercial vendors (Crl:NU(NCr)-*Foxn1nu*, Charles River Laboratories, Wilmington, MA; Hsd:Athymic Nude-*Foxn1nu(/Foxn1+)*, Inotiv, Livermore, CA (used to generate the axenic stock) and Denver, PA (used only as microbiome donors); and J:NU, The Jackson Laboratory, Bar Harbor, ME).

All mice were free of Sendai virus, pneumonia virus of mice, mouse hepatitis virus, minute virus of mice, mouse parvovirus, Theiler meningoencephalitis virus, reovirus type 3, epizootic diarrhea of infant mice (mouse rotavirus), murine adenovirus, polyoma virus, K virus, murine cytomegalovirus, mouse thymic virus, lymphocytic choriomeningitis virus, Hantaan virus, ectromelia virus, lactate dehydrogenase elevating virus, *Bordetella bronchiseptica, Citrobacter rodentium, Clostridium piliforme, Corynebacterium bovis, Corynebacterium kutscheri, Filobacterium rodentium, Mycoplasma pulmonis, Salmonella* spp., *Streptobacillus moniliformis, Streptococcus pneumoniae, Encephalitozoon cuniculi*, ectoparasites, endoparasites, and enteric protozoa. Mice received directly from vendors and housed in isocages were also free of mouse norovirus, rodent chaphamaparvovirus (except Inotiv), mouse thymic virus, *Helicobacter* spp, *Klebsiella pneumoniae, Klebsiella oxytoca, Pasteurella multocida, Rodentibacter pneumotropicus, Rodentibacter heylii, Pneumocystis murina, Proteus mirabilis, Pseudomonas aeruginosa, Staphylococcus aureus*, and beta-hemolytic *Streptococcus* spp. Based on health monitoring reports for the Inotiv PA site, mice from this location were not confirmed free of murine adenovirus, polyoma virus, Hantaan virus, ectromelia virus, lactate dehydrogenase elevating virus, mouse kidney parvovirus, mouse thymic virus, *B. bronchiseptica, F. rodentium, S. moniliformis, E. cuniculi*, and *P. multocida*.

Each stock was bred to generate timed pregnancies using copulatory plugs and weighed weekly to assess pregnancy. Litters were rederived to the germfree state via cesarean section to establish 3 distinct axenic stocks. Axenic breeding pairs (F0) were established for each stock to generate experimental mice (F1). Equal numbers of male and female 6-to 8-week-old mice were used for all experiments except the dual-infection group in Aim 3 (4 males, 2 females) due to availability (total n=152). The germfree status of each litter was confirmed at weaning by 16S PCR (IDEXX, Westbrook, ME). Prior to inoculation, mice were confirmed free of *C. bovis* and *C. amycolatum* by aerobic culture. Age-appropriate mice were allocated at random to groups within each Aim prioritizing *C. bovis*-inoculated groups over control groups for Aim 2 and 3.

### Husbandry and housing

Prior to rederivation, mice were housed in individually ventilated, autoclaved, polysulfone cages with stainless steel wire bar lids and filter tops (no. 9; Thoren Caging Systems, Hazleton, PA) on autoclaved aspen-chip bedding (PWI Industries, Quebec, Canada) with up to 3 mice per cage. Each cage was clearly identified with the stock and vendor, and provided an enrichment bag made from Glatfelter paper containing 6 g of crinkled paper strips (EnviroPak; WF Fisher and Son, Branchburg, NJ) and a 2-in. square of pulped virgin cotton fiber (cotton square; Ancare, Bellmore, NY). Each cage (including cage bottom, bedding, wire bar lid, and water bottle) was changed weekly within a vertical-flow cage changing station (AireGard NU-S619-400, NuAire, Plymouth, MN). Mice were fed a closed-source, natural ingredient, flash-autoclaved, irradiated diet (5053; LabDiet) and provided *ad libitum* reverse osmosis acidified (pH 2.5 to 2.8 with hydrochloric acid) water in polyphenylsulfone bottles with autoclaved stainless-steel caps and sipper tubes (Tecniplast, West Chester, PA).^22^

Following rederivation, mice were housed in autoclaved, solid bottom and top, gasketed and sealed, polysulfone, positive-pressure, individually ventilated cages (Isocage, Allentown Caging Equipment, Allentown, NJ) with autoclaved aspen chip bedding (PWI Industries) with up to 6 mice per cage. Cages were clearly labeled with stock, vendor site, sex, and experimental group. HEPA-filtered air (filtration at rack and cage level) ventilated each cage at approximately 30 air changes per hour. The HEPA-filtered rack effluent vented directly into the building’s HVAC system. Mice received γ-irradiated, autoclaved feed (5KA1, LabDiet) and non-acidified, autoclaved reverse osmosis water *ad libitum*. Each cage was provided with a sterile bag of Glatfelter paper containing 6 g of crinkled paper strips (EnviroPAK®, WF Fisher and Son) for enrichment. Microbiome donors were housed in isocages as described for axenic mice to maintain their specific microbiome. Cages housing axenic mice were changed at least every 8 weeks and every 2 weeks for all others. All manipulations, including cage changes, were performed in a horizontal laminar flow hood (AireGard ES NU-340, NuAire, Plymouth, MN) using sterile gloves and aseptic technique. To minimize cross-contamination, germfree mice were handled first, followed by control mice and then all other animals, with *C. amycolatum*- and *C. bovis*-inoculated mice handled last. The animal room was ventilated with 100% fresh air at a minimum of 10 air changes hourly and maintained at 72 ± 2°F (21.5 ± 1°C), relative humidity between 30% and 70%, and a 12:12 h light-dark cycle (lights on at 0600, off at 1800). All animal use was approved by Memorial Sloan Kettering’s (MSK) IACUC and conducted in accordance with AALAS’s position statements on the Humane Care and Use of Laboratory Animals and Alleviating Pain and Distress in Laboratory Animals. MSK’s animal care and use program is AAALAC-accredited and operates in accordance with the recommendations provided in the *Guide for the Care and Use of Laboratory Animals* (8th ed.).^23^

### Caesarian rederivation

Heterozygous athymic nude female mice from each stock were bred, examined for copulatory plugs, and weighed weekly to identify pregnancies. Timed-pregnant animals received 4.5 mg SQ medroxyprogesterone acetate on gestational day 17.5 to inhibit parturition. On gestational day 19.5, an aseptic field was prepared in a horizontal laminar flow hood (AireGard ES NU-340, NuAire). Each pregnant mouse was euthanized by cervical dislocation and fully submerged for 1 minute in 0.5% hydrogen peroxide (RTU Peroxigard, Virox Technologies Inc., Ontario, CA) before placement on a sterile field for a hysterectomy. The abdomen was incised, the uterus was removed *en bloc* and submerged in chlorine dioxide disinfectant solution (Clidox [1:4:1], Pharmacal Laboratories, Naugatuck, CT) for 30 seconds. The uterus was placed on a sterile heated field, each uterine horn was incised, the pups were removed and stimulated with sterile gauze until spontaneous breathing occurred. All healthy pups were transferred aseptically to an isocage containing an axenic BALB/c foster dam and 2 of the foster dam’s pups whose age was within 4 days of the litter being fostered. The foster pups’ germfree status was confirmed at weaning via 16S PCR of fecal pellets (IDEXX).

### Establishment of unique stock/microbiome combinations by cohousing

Conventionally reared female athymic nude Stock A, B, and C mice served as vendor site-specific microbiome donors. Stock A mice were sourced from two geographically distinct sites, one of which had a microbiome containing *C. amycolatum*. Microbiome donors were confirmed negative for *C. bovis* and, where applicable, *C. amycolatum* via aerobic culture. Only female mice were used as microbiome donors to minimize intracage mouse aggression.

Axenic mice of each stock (up to 4 per cage) were each cohoused for 2 weeks with a microbiome donor mouse from each of the 4 vendor sites, allowing sufficient time for microbiome reassociation through coprophagy and grooming. On arrival and between cohousing periods, microbiome donor mice were housed in isocages and handled aseptically to maintain their unique microbiome.

#### *Corynebacterium bovis* and *Corynebacterium amycolatum* propagation and inoculation

*C. bovis* isolate 7894 and *C. amycolatum* isolate 25-1107-1 stored as frozen stocks were grown on trypticase soy agar supplemented with 5% sheep blood (BBL TSA II 5% SB; Becton Dickinson, Sparks, MD) at 37°C with 5% CO□ for 48 h. Growth curves for the isolates were previously established.^1,19^ *C. bovis* isolate 7894 originated at MSK and was shown to be the most pathogenic of 6 *C. bovis* isolates evaluated in the study.^1,3^ The *C. amycolatum* isolate originated from a *C. bovis*-negative mouse at MSK with clinical signs of CAH. Inocula were prepared from bacterial suspensions during the midlog growth phase, titrated and serially diluted in brain heart infusion broth (BHI; Becton Dickinson) supplemented with 0.1% Tween 80 (VWR Chemicals, Solon, OH) to obtain 10□ CFU (*C. bovis*) and 10□ CFU (*C. amycolatum*) ± 15% bacteria in 50 μL of media. Bacterial concentrations were estimated with a MacFarland densitometer.^1^ Sterile 2-mL polypropylene tubes (Fisher Scientific, Waltham, MA) containing the inocula were transferred to the vivarium on ice, and animals were topically inoculated within 2 hours. All animal manipulations were performed in a horizontal laminar flow hood (AireGard ES NU-340) using sterile materials and aseptic technique. After inoculation, the concentration of each inoculum was confirmed and enumerated by plate count.

Control mice, inoculated with sterile media, were handled before animals inoculated with *C. bovis*. Each cage was removed from its rack, sprayed on all sides with disinfectant (Peroxigard [1:16]; Virox Technologies) and placed in the laminar flow hood. Each cage was opened for less than 5 min. Each mouse was gently grasped at the base of the tail, slightly elevating its hindquarters, and the bacterial inoculum was applied directly to the dorsal midline using a sterile filter micropipette (P200N; Marshall Scientific, Hampton, NH). A 50-μL bacterial suspension or bacterial-free BHI broth (negative controls) was deposited at 4 to 5 sites along the skin using a sterile filtered 200-μL pipette tip (Filtered Pipet Tips; Crystalgen, Commack, NY), delivering 10 to 15 μL per site to prevent immediate runoff. Cages were reassembled and returned to the rack, and new sterile gloves were donned between each cage.

### Clinical scoring

Mice were evaluated cage-side daily throughout the duration of the study, utilizing a combination of ambient light and a flashlight for improved visibility of skin lesions. The mouse’s entire skin surface was assessed and scored using the clinical scoring system described by Mendoza et al summarized in Table 1 with reference images in Figure 2.^1^ Mean clinical scores were computed by averaging clinical scores demonstrated by all mice in each respective group by day and the area under the curve (AUC) was calculated for each mouse to allow comparison between groups. Animals were euthanized prior to 21 dpi if body condition fell below 2/5 or if clinical scores reached 5.

**Figure 2.**
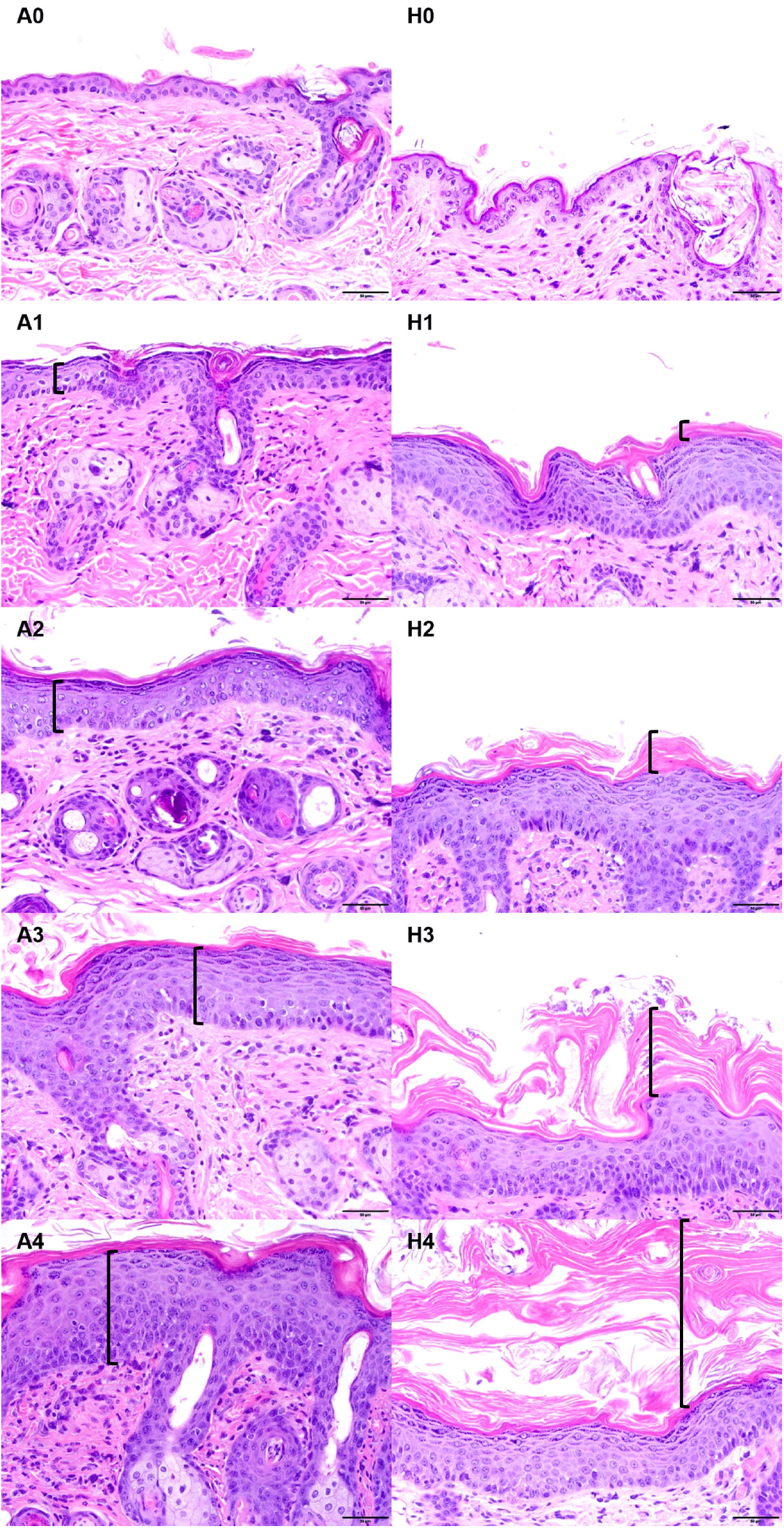
Photomicrographs highlighting the progression of acanthosis and hyperkeratosis severity in athymic nude mice infected with *C. bovis* and/or *C. amycolatum*. Samples were semiquantitatively scored as normal (0), minimal (1), mild (2), moderate (3), or severe (4) based on the intensity of acanthosis and orthokeratotic hyperkeratosis. A0 to A4, acanthosis grade; H0 to H4, hyperkeratosis grade; the black bracket represents the location and amount of acanthosis (A) or hyperkeratosis (H). Reprinted with permission from Mendoza et al.^1^

### Aerobic culture

Before inoculation and at the experimental endpoint, aerobic culture swabs were collected for *C. bovis* and, where applicable, *C. amycolatum* culture. A sterile culture swab (BBL Culture Swab, Becton Dickinson) sampled the right flank from the base of the tail cranially to the right pinna and caudally back to the tail base, rotating the swab as it was advanced.

Culture swabs were streaked in a 4-quadrant pattern onto 3 agar plates: TSA with 5% sheep blood (TSA II 5% SB; Becton Dickinson), CNA with 5% sheep blood (Columbia CNA 5% SB; Becton Dickinson), and an additional TSA plate with 5% sheep blood overlaid with a drop of Tween 80. Plates were incubated at 37°C with 5% CO□ for at least 72 hours, then evaluated for *C. bovis* and/or *C. amycolatum*, identified by small white-gray colonies characteristic for both species. Distinct colonies were speciated using MALDI-TOF spectroscopy (MALDI Biotyper Sirius CA System, Bruker; Billerica, MA). Plates without growth were held for an additional 7 days before deemed negative. Colony growth was scored from + to ++++ based on the number of quadrants containing growth.

### Postmortem gross examination and skin histopathology

On 21 dpi, mice were euthanized by CO_2_ overdose and underwent a postmortem examination, including macroscopic inspection of the skin and all internal organs. Six full-thickness skin samples, approximately 1 cm in length, were collected from the ear, head (between the ears), dorsum and ventrum (centromedial), and fore- and hindlimbs (anterior) using a scalpel. Samples were fixed in 10% neutral buffered formalin, processed in alcohol and xylene, embedded in paraffin, sectioned at 5 μm, and stained with hematoxylin and eosin. All biopsies were evaluated to confirm *C. bovis*- or *C. amycolatum-*related lesions and to quantify histologic changes. The evaluators were blinded to groups. Lesions were semi-quantitatively scored as normal (0), minimal (1), mild (2), moderate (3), or severe (4), based on acanthosis, orthokeratotic hyperkeratosis (orthokeratosis), bacterial presence, and inflammation (Figure 2), as previously described.^1^ Acanthosis, defined as increased epidermal thickness due to keratinocyte hyperplasia and often hypertrophy in the stratum spinosum, was scored as minimal (up to a 3-cell thick stratum spinosum), mild (4 to 7 cell layers), moderate (8 to 10 cell layers), and severe (>10 cell layers). Orthokeratosis, defined as increased stratum corneum thickness without retained keratinocyte nuclei and mostly identified in a compact to laminated pattern, was scored as minimal (up to 50 μm), mild (100 to 200 μm), moderate (200 to 300 μm); and severe (>300 μm). Bacterial colonies were assessed by identifying clusters of short bacilli within the stratum corneum or the lumen of hair follicles and classified as minimal (scattered bacterial rods without clusters), mild (1 to 2 clusters), moderate (3 to 5 clusters), and severe (>5 clusters). Inflammation consisted primarily of mononuclear cells and neutrophils, sometimes associated with hair follicle rupture, and was scored as minimal (single superficial pustules or scattered inflammatory cells), mild (slightly larger or more pustules or increased scattered inflammatory cells), moderate (more numerous pustules or clustering around capillaries or adnexa), and severe (large, multiple pustules with dense inflammatory cells in clusters or sheets infiltrating the dermis or adnexal epithelia). For each mouse, average scores for each of the 4 characteristics were calculated across the 6 biopsies sites and compared as individual characteristic scores and as a combined score.

### Statistical analysis

AUC and average combined histopathology scores were compared between stocks and stock-microbiome groups. Differences in AUC, mean peak clinical score, time to peak score, duration of clinical presentation, and histological scores between groups and their respective controls were compared using Kruskal-Wallis and Mann-Whitney *U* tests. AUC values represented cumulative disease burden for each mouse. Histological scores included the mean histopathology scores for combined and individual criteria for each cohort. Group size was determined to detect an effect size (Cohen’s *d*) of 2. A smaller number of control animals (n = 2 per stock per microbiome) were used for the microbiome reassociation groups to confirm that microbiome transfer alone did not induce clinical disease; these controls were excluded from statistical comparisons. A *P* value less than or equal to 0.05 was considered statistically significant. All analyses were performed using Prism 10 for Windows (GraphPad software version 10.3.0, San Diego, CA).

## Results

### Clinical scores and disease progression

#### Aim 1: C. bovis monoinoculation

The temporal course of clinical disease for axenic stocks A, B, and C monoinoculated with pathogenic *C. bovis* 7894 are shown in Figure 3. All monoinoculated mice regardless of stock developed significant clinical disease as compared to their respective sham inoculated controls (*P* ≤ 0.01), which with a single exception failed to develop lesions. Clinical disease severity and timing were most similar in Stocks A and C. In comparison, Stock B mice had a significantly lower mean AUC score, resulting from delayed onset and earlier resolution of clinical signs (*P* ≤ 0.05), as well as a lower mean peak clinical score (*P* ≤ 0.01). Sterile media inoculated controls had a clinical score of zero at all time points except a single mouse from Stock C on days 14-21 which had a score of 1.

**Figure 3.**
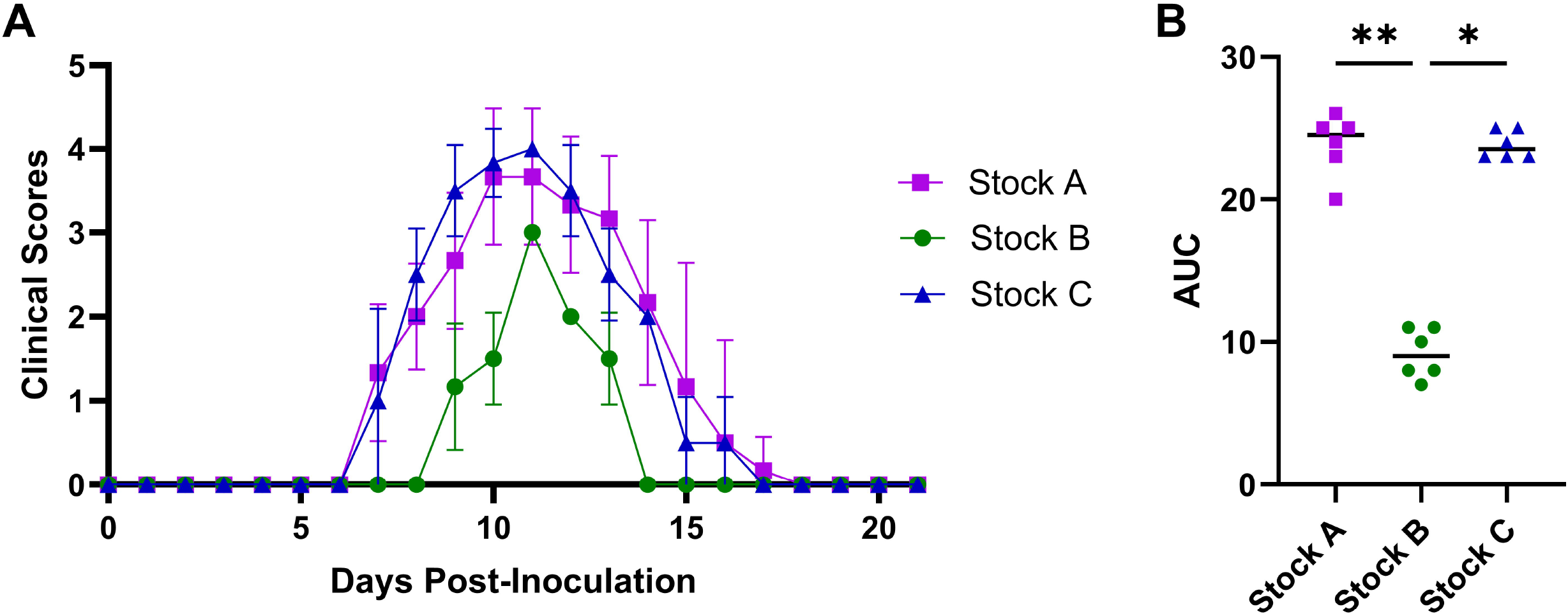
Daily Clinical Scores and Cumulative Disease Burden in Mice Monoinoculated with Pathogenic *C. bovis* 7894. A. Temporal distribution of daily disease scores (mean ± SD) for 3 outbred athymic nude mouse stocks (n = 6 mice/stock). Clinical signs were scored 0–5 based on disease severity. B. The mean area under the curve (AUC), reflecting the cumulative disease scores per stock, for the mice presented in A. Individual mice are identified by a symbol and the mean with a horizontal bar. Significant differences between stocks A and B, and B and C are shown by horizontal bars at the top of the figure. Sterile media inoculated controls are not shown. *, *P* ≤ 0.05; **, *P* ≤ 0.01.

#### Aim 2: Microbiome reassociation

Axenic mice from each of the 3 stocks were reassociated with 1 of 4 microbiomes, generating 12 unique stock/microbiome combinations, each of which was challenged with *C. bovis* 7894. The temporal distribution and cumulative disease scores for these stock/microbiome combinations are shown in Figure 4.

**Figure 4.**
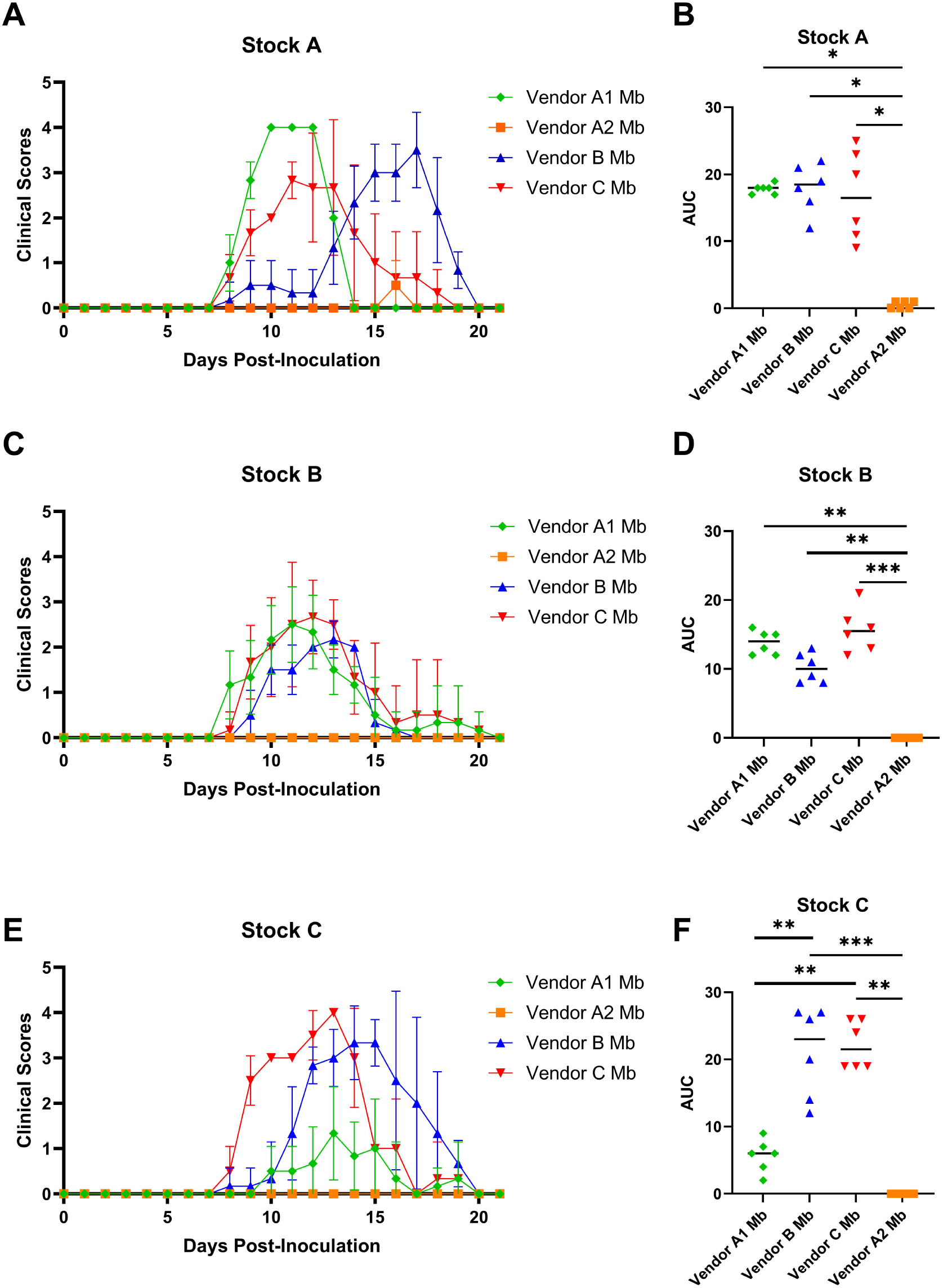
Clinical Scores and Cumulative Disease Burden for Stocks A, B, and C Reassociated with 1 of 4 Vendor Microbiomes and Challenged with *C. bovis* 7894. A, C and E: Daily mean clinical scores (± SD) for Stock A, B, and C mice (n = 6/group) reassociated with microbiomes derived from vendors A1, A2, B, and C inoculated with *C. bovis 7894*. Clinical signs were scored 0–5 based on disease severity. B, D, and F: Cumulative disease burden quantified as area under the curve (AUC) for clinical scores within each stock (n = 6/group). Individual mice are identified by a symbol and the mean with a horizontal bar. Significant differences between microbiomes by stock are shown by horizontal bars at the top of the figure. *, *P* ≤ 0.05; **, *P* ≤ 0.01; ***, *P* ≤ 0.001; Mb, microbiome.

All mice, independent of stock, reassociated with microbiomes from vendor A1, B, or C developed significant clinical lesions following inoculation with *C. bovis* 7894 (Fig. 4A–F). Importantly, none of the stocks reassociated with Vendor A2’s microbiome developed clinical disease during the study; clinical scores remained at or near zero throughout the observation period and AUC values were negligible.

Differences in temporal distribution and score severity were evident across groups. Notable was the finding that Stock B, independent of the microbiome with which it was reassociated (except A2), developed lower peak clinical scores as compared to the peak score observed with select microbiomes in Stock A and C (Fig. 4C). Across stocks, after *C. bovis* challenge, groups reassociated with Vendor B and C’s microbiome generally exhibited severe disease based on both the peak clinical scores and AUC values (Fig. 4A-F). In contrast while reassociation with Vendor A1’s microbiome resulted in severe clinical disease in Stock A and B, it caused less severe disease in Stock C. Although the microbiome was the dominant modulator of disease severity, stock-specific responses were also evident. Stock C generally exhibited higher peak scores and greater AUC values with susceptible microbiomes (Fig. 4E and F), while Stock B developed more moderate, but also more consistent disease scores (Fig. 4C and D). Stock A’s response was intermediate between the 2 other stocks (Fig. 4A and B).

The microbiome also influenced the temporal pattern of disease onset and peak after *C. bovis* challenge. Mice with Vendor A1’s microbiome generally had earlier onset but clinical disease of shorter duration in some stocks (Fig 4A, C and E). Mice with Vendor B’s microbiome generally had later disease onset as well as a more protracted disease course and mice with Vendor C’s microbiome yielded higher peak clinical disease (Fig 4A, C and E). Select microbiome effects appeared to be stock-dependent, suggesting an interaction between host genetics and the microbiome. For example, mice with Vendor A1’s microbiome had relatively lower clinical scores in Stock C (Fig. 4E), but higher clinical scores in Stocks A and B (Fig. 4A and C).

Collectively, these findings demonstrate that microbiome composition is a major determinant of susceptibility to *C. bovis-*associated disease and that specific microbial communities can confer robust protection across multiple host genetic backgrounds. Moreover, among disease-permissive microbiomes, host genetic background contributes additional variability in disease severity, indicating an important interaction between host genotype and microbiome composition in determining disease outcome.

#### Aim 3: Assessment of C. amycolatum’s pathogenicity and protection

To further investigate whether the commensal *C. amycolatum* can influence susceptibility to *C. bovis* associated disease, Stock A mice were inoculated with *C. amycolatum* alone or sequentially infected with *C. amycolatum* followed by *C. bovis* 7894 (Figure 5). Stock A mice were utilized, as they displayed significant clinical disease after inoculation with *C. bovis* 7894 in Aims 1 and 2. Axenic mice monoinoculated with *C. amycolatum* exhibited no clinical lesions throughout the study period and showed negligible disease burden as measured by the AUC, indicating that *C. amycolatum* does not cause clinical disease in this model (Fig. 5A). In contrast, mice inoculated with *C. bovis* 7894 developed characteristic clinical disease. Mice that were first colonized with *C. amycolatum* and subsequently inoculated with *C. bovis* 7894 exhibited a significant delay in the onset of clinical signs and a delayed time to peak disease severity compared with mice monoinoculated with *C. bovis* 7894 (Fig. 5A). Despite this delay in disease onset, cumulative disease burden (AUC) was comparable between the dual-infection and *C. bovis* 7894 monoinoculation groups (Fig. 5B).

**Figure 5.**
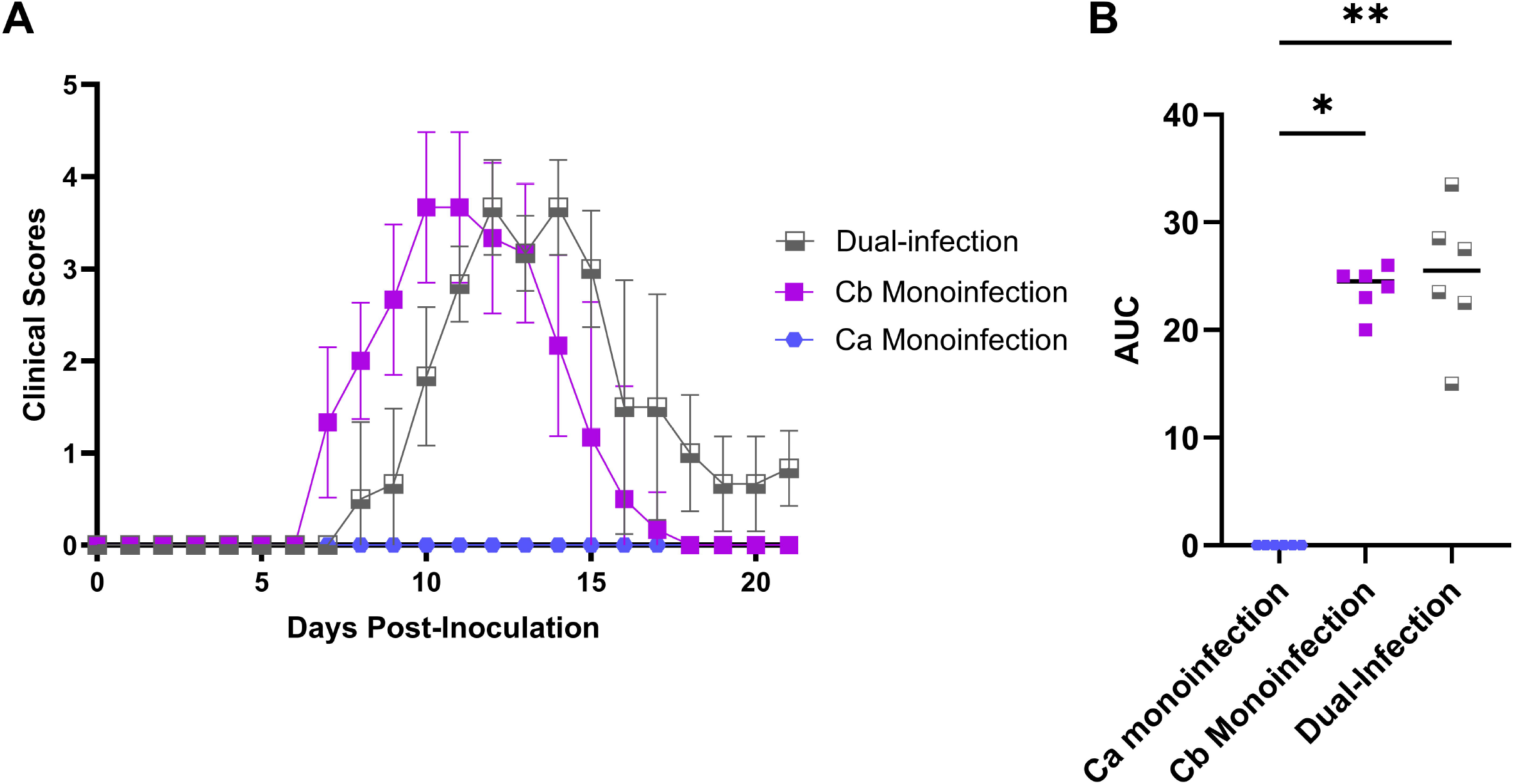
Clinical Scores and Cumulative Disease Burden in Stock A Mice Following Monoinoculation and Dual-Infection with *C. amycolatum and/or C. bovis*. A. Temporal distribution of daily disease scores (mean ± SD) for Stock A mice monoinoculated with *C. amycolatum, C. bovis* or both (n = 6 mice/group). B. The mean area under the curve (AUC), reflecting the cumulative disease scores per group for the mice presented in A. Individual mice are identified by a symbol and the mean with a horizontal bar. Significant differences between groups are shown. Day 0 post-inoculation for the dual-infection group is the day of *C. bovis* 7894 inoculation. There was no clinical disease associated with *C. amycolatum* inoculation on days –7 to –1. *, *P* ≤ 0.05; **, *P* ≤ 0.01.

Given the observed delay in disease onset associated with *C. amycolatum* colonization, we next examined whether introduction of *C. amycolatum* into a disease-permissive microbiome could alter susceptibility to *C. bovis* infection. Stock A mice were reassociated with either the A1 microbiome, the A1 microbiome supplemented with *C. amycolatum* (A1 + Ca), or the protective A2 microbiome prior to *C. amycolatum* inoculation (Figure 6). Mice harboring the A1 microbiome developed clinical disease consistent with previous observations. In contrast, mice harboring the A1 + Ca microbiome exhibited a significant delay in clinical onset (*P* ≤ 0.001) and reduced peak disease scores (*P* ≤ 0.05) compared with mice harboring the A1 microbiome alone (Fig. 6). Despite these reductions in disease severity, the protective effect of *C. amycolatum* supplementation was partial relative to mice harboring the A2 microbiome, which exhibited minimal clinical disease.

**Figure 6.**
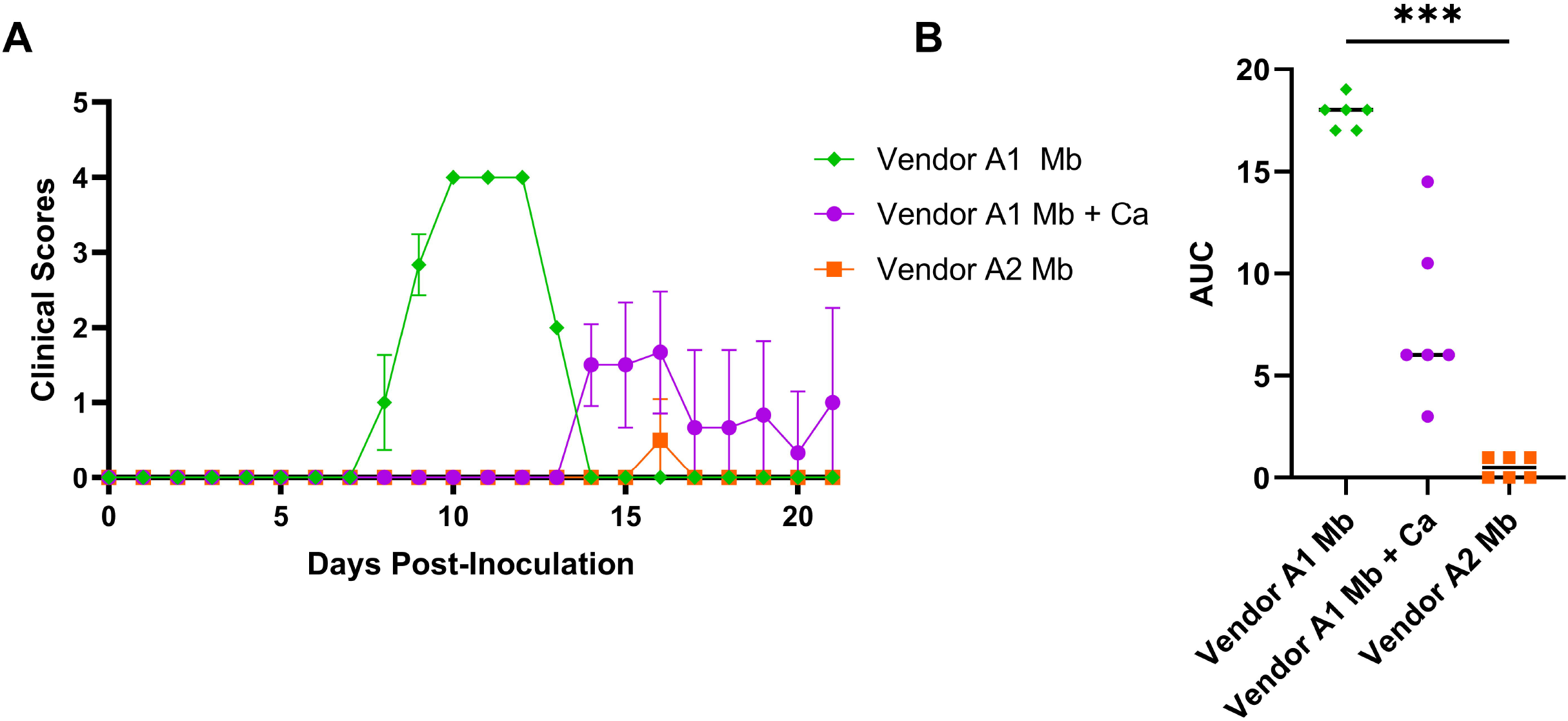
Clinical Scores and Cumulative Disease Burden in Stock A Mice Reassociated with 1 of 3 Vendor A Microbiome Variations. Stock A mice were reassociated with either the A1, A1 + *C. amycolatum*, or the A2 microbiome (n = 6/group) followed by inoculation with *C. bovis* 7894. A. Temporal distribution of daily disease scores (mean ± SD). B. The mean area under the curve (AUC), reflecting the cumulative disease scores per group for the mice presented in A. Individual mice are identified by a symbol and the mean with a horizontal bar. Significant differences between groups are shown by a horizontal bar at the top of the figure. The results for Stock A reassociated with Vendor A1 and A2 microbiomes are included from Fig. 4 for comparison. *, *P* ≤ 0.05; **, *P* ≤ 0.01; AUC, Area Under the Curve; Mb, microbiome; Ca, *C. amycolatum*.

These findings demonstrate that *C. amycolatum* does not act as a pathogen in this model but can modulate the course of *C. bovis* infection. Precolonization with *C. amycolatum* delayed disease onset and reduced peak clinical severity, and supplementation of a permissive microbiome with *C. amycolatum* similarly attenuated disease progression. These results support a role for the commensal *C. amycolatum* in microbiome-mediated modulation of susceptibility to *C. bovis* associated disease.

### Histopathology scores

#### Aim 1: C. bovis monoinoculation

To characterize the tissue-level pathology associated with *C. bovis* 7894 infection, histopathologic evaluation of skin lesions was performed in axenic Stocks A, B, and C mice following monoinoculation with *C. bovis* 7894 or sterile media. Histopathology scores were assigned individual criteria, including hyperkeratosis, acanthosis, inflammation, and bacterial colonies, and then combined to generate a cumulative score (Figure 7).

**Figure 7.**
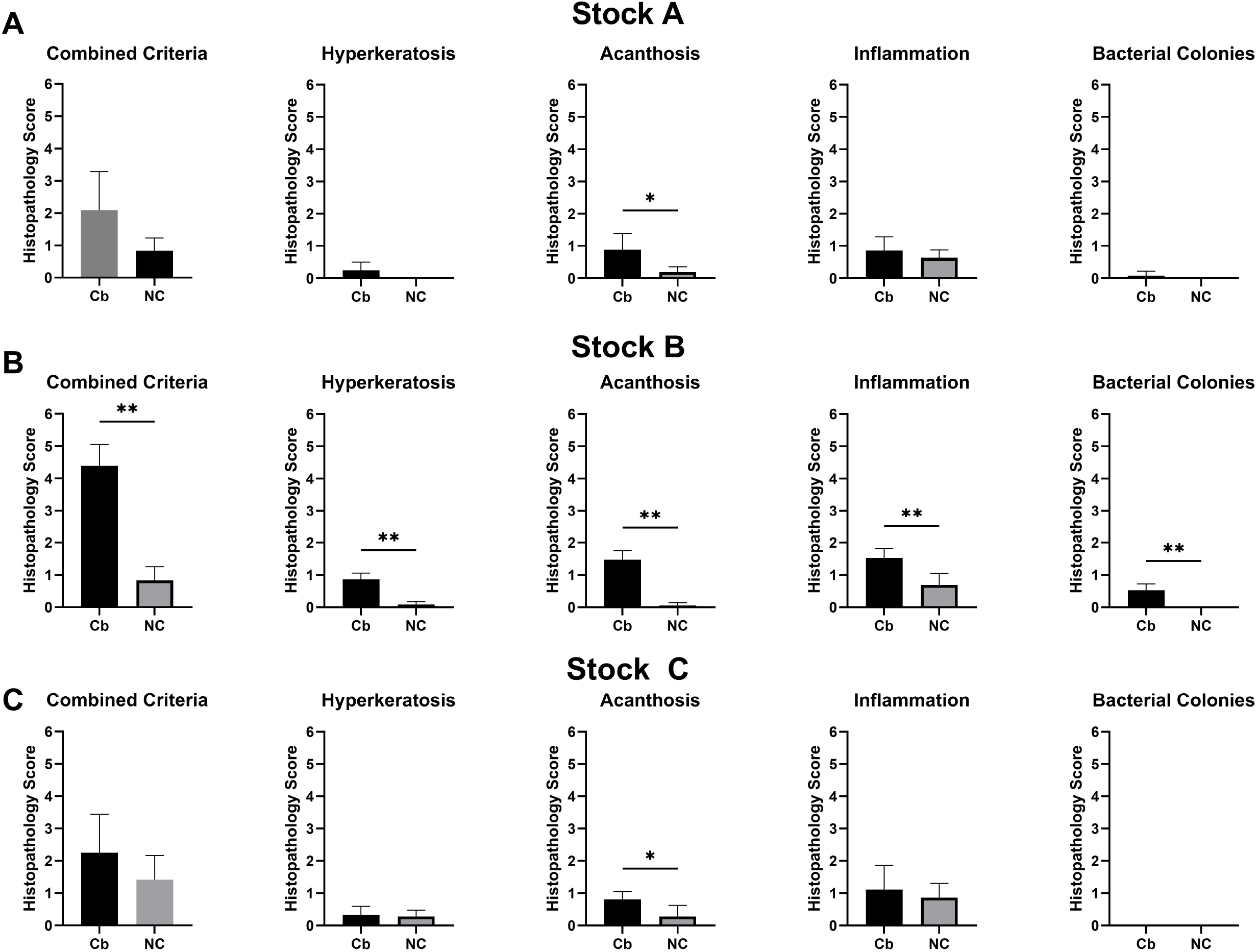
Histopathology Scores for Axenic Stock A, B, and C Mice Monoinoculated with *C. bovis* 7894. Individual criteria and the combined scores for each stock (n=6/stock; A, B, and C) at study endpoint following *C. bovis* 7894 monoinoculation are shown. Scores reflect the mean ± SD. Significant differences in scores between *C. bovis*-inoculated (Cb) and the sterile media inoculated negative control (NC) groups for each stock are reflected by a horizontal bar at the top of the figure. *, *P* ≤ 0.05; **, *P* ≤ 0.01.

Histopathologic lesions were generally mild in Stock A mice. Among the individual scoring parameters, only acanthosis scores were significantly increased in mice inoculated with *C. bovis* 7894 as compared with negative controls (*P* ≤ 0.05) (Fig. 7A). Other parameters, including hyperkeratosis and inflammation, showed modest increases but did not reach statistical significance.

In contrast, Stock B mice exhibited substantially greater histopathologic changes following *C. bovis* infection. Mice inoculated with *C. bovis* demonstrated significantly higher scores for the combined pathology score as well as for each individual criterion, including hyperkeratosis, acanthosis, inflammation, and bacterial colonies, compared with negative control animals (*P* ≤ 0.01) (Fig. 7B).

Stock C mice displayed an intermediate phenotype. Like Stock A mice, only acanthosis scores were significantly increased in *C. bovis*-inoculated animals compared with controls (*P* ≤ 0.05) (Fig. 7C), while other histopathologic parameters showed increases that did not reach statistical significance.

Across all three mouse stocks, acanthosis was the most consistently observed lesion associated with *C. bovis* 7894 infection. Acanthosis scores were significantly increased in Stock B mice in comparison with Stock C (*P* ≤ 0.05). Of the three stocks, cumulative histopathologic scores were significantly higher in Stock B mice compared to Stock A and C mice (*P* ≤ 0.05). Collectively, these findings demonstrate that *C. bovis* infection produces measurable epidermal pathology and that the magnitude of these lesions varies among mouse stocks, consistent with differences in host susceptibility observed in clinical disease scoring.

#### Aim 2: Microbiome reassociation

Histopathology scores were assessed across mouse stocks (A, B, and C) reassociated with each of the four distinct vendor microbiomes (A1, A2, B, and C) to evaluate the impact of host genetics and microbiome composition on tissue-level pathology associated with *C. bovis* 7894 infection. Figure 8 provides both the combined and individual scores for each of the 12 combinations.

**Figure 8.**
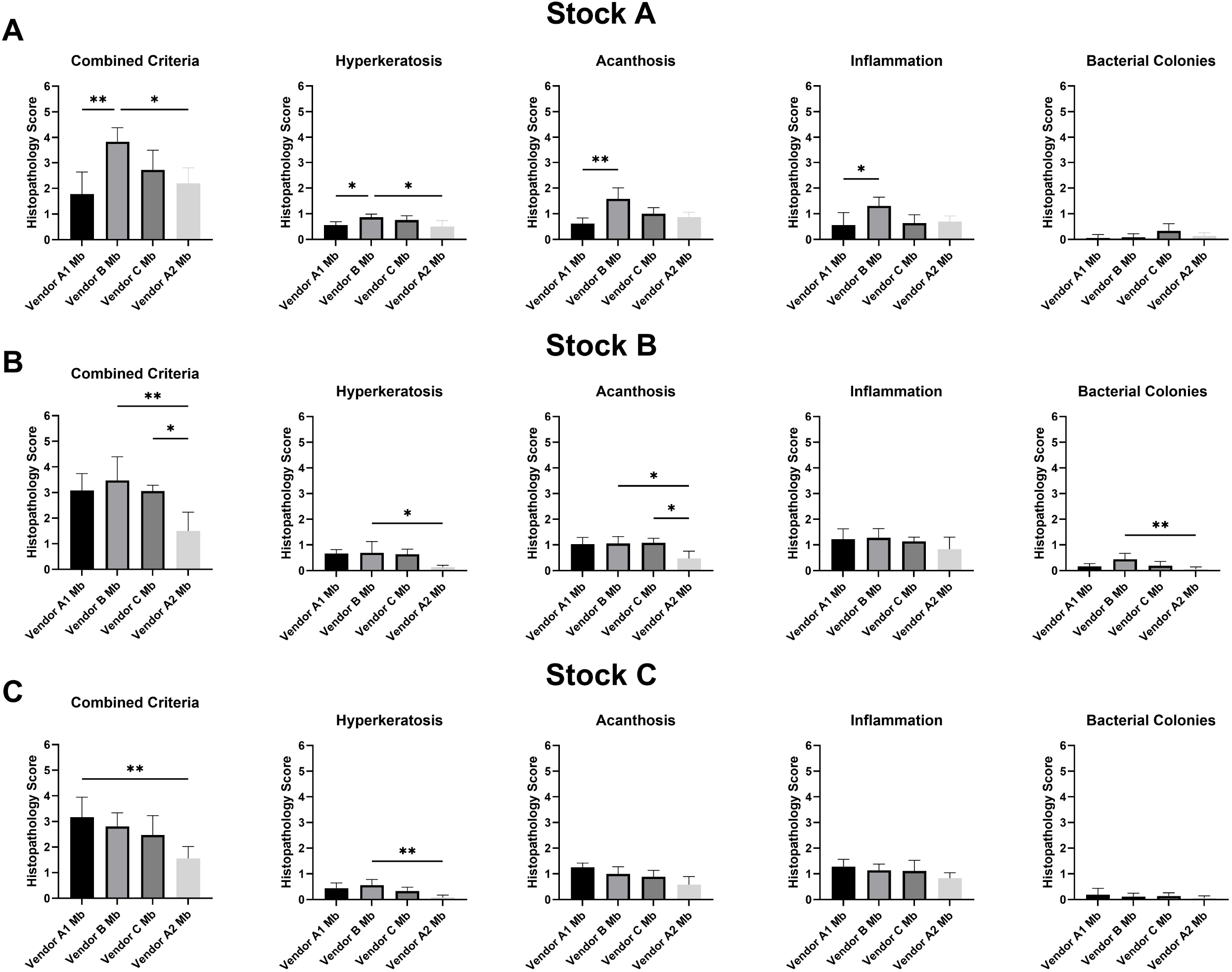
Histopathology Scores for Each Stock Reassociated with 1 of 4 Microbiomes Followed by *C. bovis* 7894 Inoculation. Individual and combined histopathology scores for Stock A (A), Stock B (B), and Stock C (C) mice reassociated with each of the 4 microbiomes (Vendors A1, A2, B, and C) inoculated with *C. bovis* 7894 (n = 6/group). Significant differences in scores between microbiomes for each stock are reflected by a horizontal bar at the top of the figure. *, *P* ≤ 0.05. **, *P* ≤ 0.01.

In Stock A, mice harboring the Vendor B microbiome exhibited significantly higher combined histopathology scores than mice harboring the Vendor A1 (*P* ≤ 0.01) or Vendor A2 microbiomes (*P* ≤ 0.05) (Fig. 8A). Hyperkeratosis scores were significantly greater in mice harboring the Vendor B and C microbiomes compared with the Vendor A2 microbiome (*P* ≤ 0.05). Acanthosis scores were significantly elevated in mice harboring the Vendor B microbiome compared with Vendor A1 (*P* ≤ 0.01), and inflammation scores were higher in mice harboring the Vendor B microbiome compared with Vendor A1 (*P* ≤ 0.05).

In Stock B, combined histopathology scores were significantly higher in mice harboring the Vendor B microbiome compared with the Vendor A2 microbiome (*P* ≤ 0.01), while mice harboring the Vendor C microbiome also exhibited higher combined scores than the Vendor A2 group (*P* ≤ 0.05) (Fig. 8B). Hyperkeratosis scores significantly increased in mice harboring the Vendor C microbiome relative to Vendor A2 (*P* ≤ 0.05). Acanthosis scores were significantly higher in mice harboring Vendor B and Vendor C microbiomes compared with Vendor A2 (*P* ≤ 0.05), and bacterial colony scores were greater in mice harboring the Vendor B microbiome than in mice harboring the Vendor A2 microbiome (*P* ≤ 0.01).

In Stock C, mice harboring the Vendor A2 microbiome exhibited significantly lower combined histopathology scores compared with mice harboring the Vendor A1 microbiome (*P* ≤ 0.01) (Fig. 8C). Hyperkeratosis scores were also significantly reduced in the Vendor A2 group compared with the Vendor C microbiome group (*P* ≤ 0.01). No significant differences were observed among microbiome groups for acanthosis, inflammation, or bacterial colony scores.

Collectively, these results demonstrate that microbiome composition significantly influences the histopathologic severity of *C. bovis-*associated disease, with the Vendor B microbiome generally associated with greater lesion severity and the Vendor A2 microbiome associated with reduced pathology, consistent with the clinical disease patterns observed in earlier experiments.

#### Aim 3: Assessment of C. amycolatum’s pathogenicity and protection

Histopathologic lesion severity differed among groups monoinoculated with either *C. bovis, C. amycolatum* or both (Figure 9). Combined pathology scores were significantly higher in the *C. bovis* and dual-infection groups than in the *C. amycolatum* group (*P* ≤ 0.05). Hyperkeratosis scores were significantly increased in the dual-infection group relative to negative controls (*P* ≤ 0.001). Acanthosis scores were significantly higher in both the *C. bovis* and dual-infection groups than in the *C. amycolatum* and control groups (*P* ≤ 0.01). Inflammation and bacterial colony scores were low and did not differ significantly among groups. Collectively, these findings indicate that cutaneous pathology was associated with the presence of *C. bovis*, whereas *C. amycolatum* alone produced minimal lesions comparable to axenic controls.

**Figure 9.**
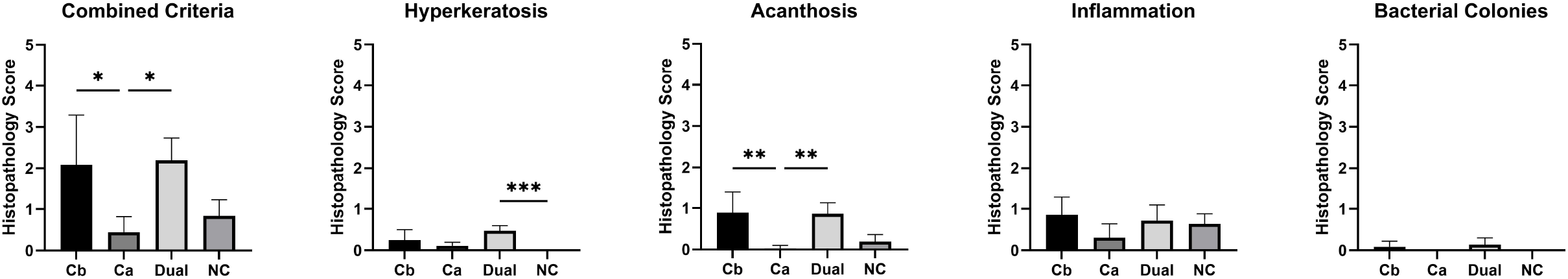
Histopathology Scores for Stock A Monoinoculated with *C. bovis* 7894, *C. amycolatum* or Both. Individual and combined histopathology scores for axenic Stock A mice inoculated with *C. bovis* 7894, *C. amycolatum* both or sterile media (n = 6/group). Scores reflect the mean ± SD. Significant differences in scores between groups are reflected by a horizontal bar at the top of the figure. *, *P* ≤ 0.05; **, *P* ≤ 0.01; ***, *P* ≤ 0.001; Dual, inoculated with both bacterial species; NC, negative sterile media inoculated control.

Histopathologic lesion severity was compared in Stock A mice reassociated with the nonprotective Vendor A1 microbiome, the protective A2 microbiome which includes *C. amycolatum*, and the A1 microbiome supplemented with *C. amycolatum* inoculated with *C. bovis* 7894 (Figure 10). The Vendor A1 microbiome supplemented with *C. amycolatum* group exhibited significantly higher combined pathology scores than the Vendor A1 (*P* ≤ 0.01) and the Vendor A2 (*P* ≤ 0.05) microbiome groups. Similar patterns were observed for individual criteria: hyperkeratosis scores were significantly higher in the Vendor A1 microbiome supplemented with *C. amycolatum* group than in both comparison groups (*P* ≤ 0.05), acanthosis scores were significantly increased relative to the Vendor A1 microbiome group (*P* ≤ 0.01), and inflammation scores were significantly higher than those of the Vendor A1 microbiome (*P* ≤ 0.01) and Vendor A2 microbiome (*P* ≤ 0.05) groups. Bacterial colony scores remained low across all groups and did not differ significantly. Clinical scores for this group were low, but persisted through the study’s endpoint. Together, these results indicate persistent skin pathology in mice harboring the Vendor A1 microbiome after *C. amycolatum* was introduced.

**Figure 10.**
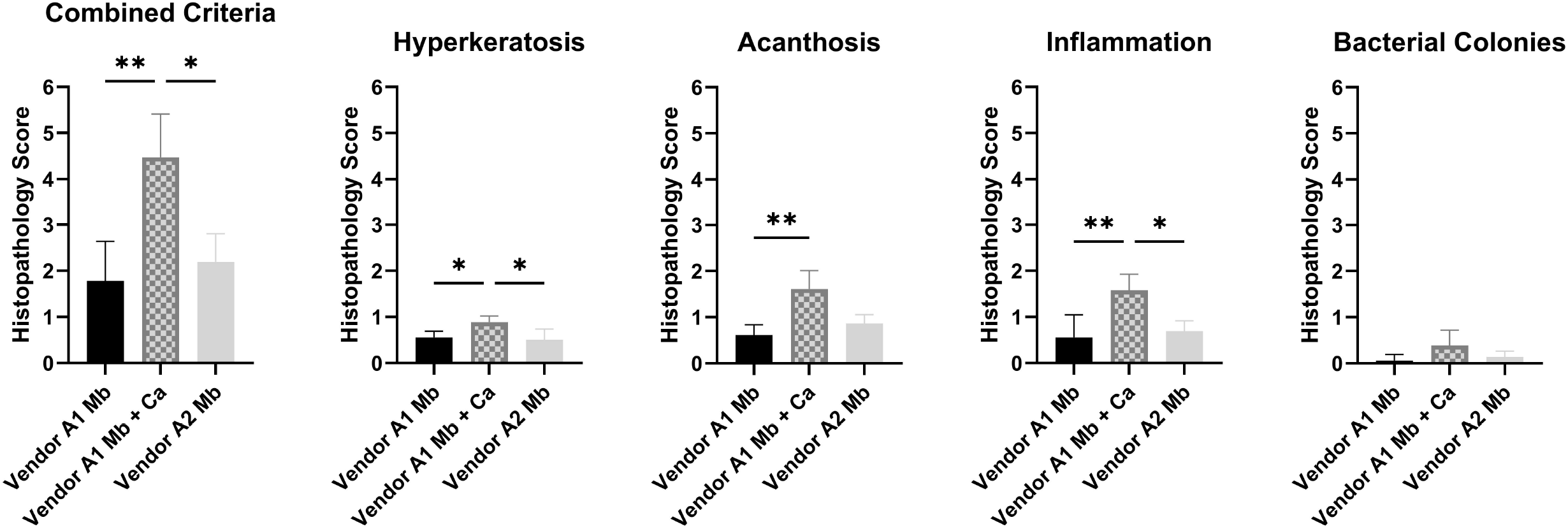
Histopathology Scores for Stock A Mice Reassociated with 3 Distinct Microbiomes Challenged with *C. bovis* 7894. Microbiome reassociated groups (n=6/group) included Vendor A1, Vendor A1 supplemented with *C. amycolatum*, and Vendor A2. Significant differences in scores between groups are reflected by a horizontal bar at the top of the figure. *, *P* ≤ 0.05; **, *P* ≤ 0.01; Mb, microbiome; Ca, *C. amycolatum*.

## Discussion

The present study demonstrates that infection with *C. bovis* can induce *Corynebacterium*-associated hyperkeratosis (CAH) in axenic outbred athymic nude mice and that both host genetic background and microbiome composition influence the severity and temporal progression of disease. These findings support the concept that CAH severity represents the outcome of interactions between the bacterium and the resident microbiome modified by host genetics.

Monoinfection experiments demonstrated that *C. bovis* is sufficient to induce CAH in nude mice. Axenic mice from all 3 stocks developed characteristic clinical lesions following topical inoculation with a defined preparation of *C. bovis* 7894, and the organism was recovered from affected animals at the experimental endpoint. This *C. bovis* isolate was previously isolated from naturally infected mice and shown to reproducibly induce disease in experimental studies.^1,2,19^ The use of axenic mice establishes that *C. bovis* alone is capable of causing CAH in the absence of other microbial influences, thereby excluding a requirement for microbiome-derived cofactors in disease induction. Taken together with prior work,^1,2,8,19^ these findings satisfy Koch’s postulates and demonstrate that *C. bovis* is the etiologic agent of CAH in the athymic nude mouse.^24,25^

Although clinical disease was observed in all 3 stocks, disease severity differed among host genetic backgrounds. Stock B mice exhibited delayed clinical disease onset, lower peak clinical scores, and reduced cumulative disease burden as compared with Stocks A and C. These findings demonstrate for the first time that genetic differences can influence susceptibility to *C. bovis*-associated disease. Because all stocks share the *Foxn1* mutation responsible for athymia, these differences likely reflect additional genetic variation affecting, among other things, epidermal barrier integrity, inflammatory signaling, or host-microbe interactions. Vendor-associated differences in CAH severity have been reported previously but the study could not discriminate between role of host genetics, the microbiome or both.^2^

Histopathologic findings in the monoinfected mice also revealed different pattern differences across host genetic backgrounds. Stock B mice demonstrated significantly higher total histopathology scores compared with negative controls, whereas in Stocks A and C only acanthosis scores differed significantly from controls. Although overall pathology scores were generally low across groups, acanthosis was the most consistent lesion associated with infection, consistent with previously described epidermal hyperplasia in CAH.^1,2,4,8,26^ The discrepancy between clinical disease severity and histopathology scores among stocks may reflect differences in host responses to persistent bacterial colonization. Greater numbers of bacterial colonies were observed histologically in Stock B mice, which could perpetuate epidermal hyperplasia and low-grade inflammation even after clinical lesions resolved. Alternatively, variations may reflect differences in the timing of lesion resolution relative to the study endpoint. Epidermal hyperplasia often persists during the healing phase after clinical lesions diminish; therefore, histopathologic findings may reflect lesion chronology rather than active disease severity. Specifically, the observed changes may simply reflect the timing relative to the course of disease at which the samples were collected and analyzed; most samples were collected after clinical lesion resolution. For instance, animals whose clinical lesions resolved just prior to collection would be expected to exhibit more severe histopathologic lesions compared to those whose lesions resolved earlier, allowing more time for tissue remodeling. Discrepancies in clinical severity and histopathology scores may also reflect limitations of semi-quantitative scoring and the selection of 21 dpi as the endpoint.

One negative control Stock C mouse developed mild localized flaking without detectable *C. bovis* or other aerobic bacteria on repeated culture. The lesions were mild, confined to the ventral abdomen and hindlimbs, did not progress during the study period, and were not associated with the characteristic distribution of CAH. Such findings may represent a noninfectious inflammatory dermatosis, such as an immune-mediated interface-like dermatitis occasionally seen in athymic nude mice and rats (unpublished data), or other types of spontaneous dermatitis described in immunodeficient strains. The data from this animal were retained in the analysis and did not alter the statistical outcomes.

Microbiome reassociation experiments demonstrated that microbial community composition is a major determinant of CAH susceptibility. Mice from all 3 stocks reassociated with the Vendor A2 microbiome failed to develop clinically significant CAH following *C. bovis* challenge, whereas mice reassociated with microbiomes from Vendors A1, B, or C developed variable but frequently substantial disease. These results demonstrate that certain microbial communities can prevent the development of clinical disease even in genetically susceptible hosts. Microbiome-mediated resistance to pathogen colonization is well documented in both gastrointestinal and cutaneous systems and may occur through competitive exclusion, production of inhibitory metabolites, or modulation of host immune responses.^14-16,27^ Although the specific microbial taxa responsible for protection remain undefined, these findings demonstrate that microbial community composition can strongly influence susceptibility to CAH.

Despite the dominant role of microbiome composition, host background continued to influence clinical outcomes. Differences in disease kinetics and severity were observed among stocks harboring the same microbiome, suggesting that host genetic factors modulate interactions between commensal and pathogenic microorganisms. For example, Stock C mice reassociated with the Vendor A1 microbiome exhibited lower clinical disease severity than mice from other stocks harboring the same microbiome. These observations indicate that host genotype and microbiome composition interact to shape disease resistance, with the latter increasingly recognized in models of infectious and inflammatory skin disease.^28^

Histopathologic findings in the microbiome reassociation experiments generally paralleled clinical observations, and differences among groups were often modest. Mice harboring the Vendor A2 microbiome exhibited the lowest pathology scores, consistent with the absence of clinical lesions. Groups with higher pathology scores frequently corresponded to cohorts in which clinical lesions resolved later in the study period. This apparent discordance likely reflects differences in the timing of lesion resolution relative to necropsy, as epidermal hyperplasia and hyperkeratosis may persist during the healing phase even after overt clinical lesions have diminished. Semi-quantitative histopathology scoring is inherently subjective; however, all sections in this study were evaluated by a single individual and reviewed by a board-certified comparative pathologist utilizing established scoring criteria to minimize interobserver variability.

The role of the commensal bacterium, *C. amycolatum*, was evaluated to determine its contribution to microbiome-mediated modulation of CAH. This investigation was prompted by the observation that *C. amycolatum* was present in the skin microbiome of nude mice from only one of two Vendor A facilities. Notably, mice from this specific site developed less severe clinical disease when inoculated with *C. bovis*. Furthermore, *C. amycolatum* was identified as a microbiome constituent in an additional nude mouse stock that appeared resistant to *C. bovis*-associated disease.^2^ The isolate used in this study was originally recovered from the skin of a mouse with dermatitis but did not produce clinical disease when inoculated as a monoinfection in axenic mice. Histopathology scores in mice colonized with *C. amycolatum* alone were minimal and comparable to those observed in axenic negative controls, indicating that this organism is not independently pathogenic in this model.

Precolonization of axenic mice with *C. amycolatum* prior to *C. bovis* infection produced a modest delay in disease onset and time to peak clinical severity, although cumulative disease burden was not significantly reduced. These findings suggest that *C. amycolatum* alone is insufficient to prevent CAH but may alter early pathogen colonization dynamics or host responses in ways that temporarily limit disease progression. Interestingly, supplementation of the Vendor A1 microbiome with *C. amycolatum* produced a more pronounced alteration in disease course, including delayed onset and reduced clinical severity compared with the Vendor A1 microbiome alone. These findings suggest that *C. amycolatum* may function as a modulatory commensal organism that interacts with other members of the microbiome to influence disease outcomes rather than acting as a single protective keystone species.

A potential explanation for variability in clinical severity among infected animals is the influence of the resident skin microbiome on pathogen colonization and expansion. Commensal microbial communities are increasingly recognized as important mediators of colonization resistance, whereby resident organisms inhibit the establishment or proliferation of potential pathogens through niche occupation, metabolic competition, or modulation of host immune responses. Our prior observations that colonization with *C. amycolatum* was associated with partial attenuation of disease severity raise the possibility that competition within the cutaneous *Corynebacterium* niche may influence the clinical expression of infection with *C. bovis*. Such interactions may occur through direct ecological competition at the level of hair follicles and superficial epidermis, or indirectly through effects on keratinocyte immune signaling and antimicrobial peptide production. More broadly, these findings align with the concept that both local skin microbial communities and systemic immune signals derived from the intestinal microbiome can shape inflammatory responses at epithelial barriers. Together, these interactions suggest that variation in microbiome composition, whether at the level of the skin and/or gut, may represent an underappreciated determinant of disease phenotype in *C. bovis* infection.

Histopathology scores in the microbiome supplemented with *C. amycolatum* were higher than those observed in the initial Vendor A1 microbiome group despite reduced clinical disease severity. This apparent discrepancy likely reflects differences in lesion resolution timing relative to necropsy, as previously noted. Histologic lesions may have equalized between groups if the endpoints were based on clinical resolution rather than a defined period following inoculation. These findings emphasize that clinical and histopathologic assessments measure different aspects of disease progression and should be interpreted together when evaluating CAH models.

Several limitations should be considered when interpreting these data. This study did not address specific microbiome components other than the presence or absence of C. *amycolatum*. Future metagenomic analysis is planned. Additionally, microbiome reassociation was achieved through cohousing rather than vertical transmission from dams. Cohousing does not fully replicate natural acquisition of microbiota early in life and potentially affects early immune system development, but for the purposes of this study it allowed controlled reassociation of axenic mice with defined microbiome sources prior to experimental infection. Also, bacterial colonization was monitored primarily by culture, and molecular detection methods such as PCR could provide greater sensitivity detecting low-level colonization. Study results may also have been impacted by limited group size.

Collectively, these findings further define the clinical and histopathologic features associated with infection by *C. bovis* in athymic nude mouse stocks and highlight the potential role of microbial community structure in shaping disease expression. The observation that colonization with *C. amycolatum* may partially mitigate clinical disease supports the concept that interactions within the resident skin microbiome can influence pathogen colonization dynamics and host inflammatory responses. Furthermore, evidence that intestinal microbiota influence systemic immune homeostasis suggests that both local and extra-intestinal microbial communities may drive variations in clinical phenotypes. The insights provided in this study may also have practical implications for colony management and model development. For example, improved understanding of microbiome-pathogen interactions could potentially inform the generation of nude mouse stocks with increased resistance to *C. bovis*-associated disease. However, because historically generated datasets are often tied to specific mouse stocks and their associated microbial communities, the introduction of new stocks with altered microbiome compositions may influence experimental outcomes and complicate comparisons with prior studies. Consequently, while the development of more disease-resistant stocks may offer advantages for colony health, careful consideration must be given to the potential impact of microbiome differences on experimental reproducibility. Future studies integrating longitudinal characterization of skin and gut microbiomes with experimental colonization models will be important for clarifying the relative contributions of these microbial communities to disease susceptibility and for informing strategies to mitigate dermatologic disease in immunodeficient mouse colonies while preserving research fidelity.

## Acknowledgements

We acknowledge Callista Huang, Nimisha Pattada, Abigail Michelson, Glory Leung, Juliette Wipf, Melissa Nashat, Felix Wolf, Sarah Paisner, and the staff from the Laboratory of Comparative Pathology for assistance with various technical components of the study. Generative artificial intelligence (ChatGPT, GPT-5.3, OpenAI and Gemini 3 Flash, Google) was used to assist with drafting and improving clarity of the manuscript text. All content was reviewed and finalized by the authors.

## Data access

Datasets and research notes are available at MSK.

